# The effect of waning on antibody levels and memory B cell recall following SARS-CoV-2 infection or vaccination

**DOI:** 10.1101/2022.03.16.484099

**Authors:** David Forgacs, Vanessa S. Moraes, Hannah B. Hanley, Jasper L. Gattiker, Alexandria M. Jefferson, Ted M. Ross

**Author notes:** To whom correspondence shall be addressed: Ted M. Ross, Ph.D., Center for Vaccines and Immunology, University of Georgia, 501 D.W. Brooks Drive, CVI Room 1504, Athens, GA 30602.

## Abstract

As of March 2022, there have been over 450 million reported SARS-CoV-2 cases worldwide, and more than 4 billion people have received their primary series of a COVID-19 vaccine. In order to longitudinally track SARS-CoV-2 antibody levels in people after vaccination or infection, a large-scale COVID-19 sero-surveillance progam entitled SPARTA (SeroPrevalence and Respiratory Tract Assessment) was established early in the pandemic. Anti-RBD antibody levels were tracked in more than 1,000 people. There was no significant decrease in antibody levels during the first 14 months after infection in unvaccinated participants, however, significant waning of antibody levels was observed following vaccination, regardless of previous infection status. Moreover, participants who were pre-immune to SARS-CoV-2 prior to vaccination seroconverted to significantly higher antibody levels, and antibodies were maintained at significantly higher levels than in previously infected, unvaccinated participants. This pattern was entirely due to differences in the magnitude of the initial seroconversion event, and the rate of antibody waning was not significantly different based on the pre-immune status. Participants who received a third (booster) dose of an mRNA vaccine not only increased their anti-RBD antibody levels ∼14-fold, but they also had ∼3 times more anti-RBD antibodies compared to the peak of their antibody levels after receiving their primary vaccine series. In order to ascertain whether the presence of serum antibodies is important for long-term seroprotection, PBMCs from 13 participants who lost all detectable circulating antibodies after vaccination or infection were differentiated into memory cells *in vitro*. There was a significant recall of memory B cells in the absence of serum antibodies in 70% of the vaccinated participants, but not in any of the infected participants. Therefore, there is a strong connection between anti-RBD antibody levels and the effectiveness of memory B cell recall.

## Introduction

In late 2019, Severe Acute Respiratory Syndrome Coronavirus 2 (SARS-CoV-2), the causative agent of COVID-19, emerged in Wuhan, China, and quickly spread across the world resulting in the ongoing COVID-19 pandemic^1^. As the virus continues to evolve and adapt to humans, several viral variants of concern have emerged, and most likely, new variants will continue to emerge in the future that could result in severe human disease^2^.

Antibody-mediated immunity is essential in order to mount a systemic immune response against SARS-CoV-2. The receptor binding domain (RBD) located on the tip of the spike glycoprotein is one of the major targets of neutralizing antibodies^3,4^. The virus uses the spike protein to bind to the ACE2 receptors on the surface of host cells in order to gain cell entry^3,4^. Therefore, several COVID-19 vaccines were developed based on the spike protein as an immunogen^5,6^. The Pfizer-BioNTech BNT162b2 and the Moderna mRNA-1273 vaccines both contain mRNA coding for the full-length spike protein^5,6^. These vaccines are administered intramuscularly as two doses 21 or 28 days apart respectively^5,6^. A third, single-dose viral vector vaccine by Janssen also received emergency use authorization in the U.S.^7,8^. A individual is considered fully vaccinated 14 days after they completed the primary vaccine series^5,9^.

Anti-spike antibodies increase significantly following vaccination^10,11^. However, there is a significant drop in antibody levels within the first few months following vaccination^12–14^. In order to keep the level of circulating antibodies high, a third (booster) dose of the Pfizer-BioNTech and Moderna vaccines was approved in the U.S. during the second half of 2021^15^. While all Pfizer-BioNTech vaccine doses – including the third (booster) dose – contain 30 µg mRNA, the primary Moderna vaccine series contain 100 µg, and the booster dose contains only 50 µg^5,9,15,16^.

Besides humoral immunity, cellular immunity is also essential for long term protection against infection^17,18^. Memory B cells and long lived plasma cells created due to infection or vaccination can be recalled in the event of a repeat encounter, yielding a faster and more robust antibody response^17^. The relationship between the levels of circulating and memory recall response is an underexplored area of SARS-CoV-2 immunology research.

In 2020, SPARTA (SARS SeroPrevalence and Respiratory Tract Assessment), a study funded by the U.S. National Institutes of Health (NIH) was initiated to understand if immunity elicited following infection with SARS-CoV-2 or vaccination with COVID-19 vaccines provides protection against future infection and symptomatic disease^10^. The goals were 1) to investigate the level and duration of protection afforded by natural infection following SARS-CoV-2 infection, 2) to assess immunological risk factors for infection outcome and examine immune responses to infection across the disease spectrum, and 3) to study the immune effectiveness of COVID-19 vaccines in both pre-immune and immunologically naïve participants. Participants were located at several sites in three U.S. states and represented a diverse cohort that provided a real-time snapshot into the progression of immune responses during the pandemic. To date, ∼3,800 participants have provided serum, peripheral blood cells, and saliva at multiple timepoints.

One of the goals of this study was to examine the rate of antibody waning in a large cohort of people infected with SARS-CoV-2 and/or vaccinated with a commercial COVID-19 vaccine, as well as to discover differences in trends associated with antibody levels based on vaccination, infection, and pre-immunity. In addition, in order to ascertain whether the presence of serum antibodies is necessary for a robust recall response, peripheral blood mononuclear cells (PBMCs) were collected and differentiated into memory B cell *in vitro* from 13 individuals. These 13 participants had an initial antibody response induced by vaccination or infection, but their antibody levels subsequently declined to undetectable levels. In the absence of serological protection, the *de novo* memory B cell recall response was analyzed in these participants.

## Materials and Methods

### SPARTA participants

Eligible participants between 18 and 90 years old were enrolled starting in April 2020 with written informed consent in Athens and Augusta, GA, Memphis, TN, and Los Angeles, CA. The study procedures, informed consent, and data collection documents were reviewed and approved by the WIRB Copernicus Group Institutional Review Board (WCG IRB #202029060). Of the ∼3,800 SPARTA enrolled participants, 1,081 were randomly selected to be included in this study (Table S1). 68.5% of them identified as females, and 31.4% as males. The average age was 44.9 years (median age = 44 years old, SD = 17.2 years). 86.1% of the cohort identified as White, 7.3% as Black/African American, 4.1% as Asian, and 1.6% as multiple races. 8.1% of the participants were Hispanic. The range of BMI was between 17.5 and 95.5, with an average BMI of 28.5 (median BMI = 27.2, SD = 6.8).

### Enzyme-linked immunosorbent assay (ELISA)

ELISA assays were performed as previously described^10^. Briefly, Immulon^®^ 4HBX (Thermo Fisher Scientific, Waltham, MA, USA) or Costar EIA/RIA (Corning, Corning, NY, USA) plates were coated with 100 ng/well of recombinant SARS-CoV-2 RBD protein, incubated with heat inactivated serum samples at a starting dilution of 1:50 and then further serially diluted 3-fold^19^. IgG antibodies were detected using horseradish peroxidase (HRP)-conjugated goat anti-human IgG detection antibody (Southern Biotech, Birmingham, AL, USA) at a 1:4,000 dilution and colorimetric development was accomplished using 100 μL of 0.1% 2,2’-azino-bis(3-ethylbenzothiazoline-6-sulphonic acid) (ABTS, Bioworld, Dublin, OH, USA) solution with 0.05% H_2_O_2_ for 18 minutes at 37ºC. The reaction was terminated with 50μL of 1% (w/v) SDS (VWR International, Radnor, PA, USA). Colorimetric absorbance was measured at 414nm using a PowerWaveXS plate reader (Biotek, Winooski, VT, USA). All samples and controls were run in duplicate, and the mean of the two blank-adjusted optical density (OD) values were used in downstream analyses. IgG equivalent concentrations were calculated based on a 7-point standard curve generated by a human IgG reference protein from plasma (Athens Research and Technology, Athens, GA, USA), and verified on each plate using human sera of known concentrations.

### Viral neutralization assay

Viral neutralization (VN) assays were performed in a Biosafety Level 3 (BSL-3) laboratory as previously described^10^. The USA-WA1/2020 SARS-CoV-2 strain (100 TCID50/50µl; NCBI accession number: PRJNA717311) was co-incubated with serially diluted serum samples for 1 hour at 37°C and then added to a monolayer of Vero E6 cells. The plates were observed after 72 hours for cytopathic effects (CPEs). The VN endpoint titer was determined as the reciprocal of the highest dilution that completely inhibited CPE formation. All neutralization titers were represented as the average of the three replicates.

### In vitro differentiation

PBMCs were differentiated into memory B cells by incubating them with 500 ng/mL R848 (Invivogen, San Diego, CA, USA) and 5 ng/mL rIL-2 (R&D, Minneapolis, MN, USA) for 7 days at 37ºC in 5% CO_2_ as previously described^20^. The conditioned cell culture supernatants were serially diluted to assess total and antigen-specific IgG antibody levels by ELISA. Total (non-antigen-specific) IgG levels were assessed to confirm that the *in vitro* differentiation was successful and B cells were actually recalled *de novo*. Antigen-specific IgG levels to SARS-CoV-2 RBD, and full-length spike protein, as well as pandemic influenza strain A/H1N1/California/2009 (Cal/09) were also measured, along with spike-specific IgA and IgM levels.

### Direct ex vivo B cell immune cell phenotyping

Cryopreserved cells were thawed by diluting them in 10 mL pre-warmed complete B cell media (RPMI + 5% FBS + 1% Pen/Strep) in the presence of DNAse (20 µg/mL) and spun at 500 *x* g for 10 min. Supernatants were carefully removed, and the cells were counted and rested at 2×10^6^ cells/mL for 3 hours. After resting, cells were washed again in complete B cell media and 1×10^6^ cells were resuspended in 50 µl of PBS containing LIVE/DEAD (Thermo Scientific) and human FC block (1:50, Biolegend), and incubated for 20 min at room temperature (RT). After incubation, cells were topped with 150 µl of PBS with 2% FBS (FACS buffer) and spun 500 *x* g for 4 min. The supernatant was removed and the antibody mix containing the surface antibodies was added to the cells and incubated for 30 min at RT in the dark (Figure S1). Following surface staining, cells were washed twice with FACS buffer and resuspended in a final volume of 200 µL. Samples were acquired on Agilent NovoCyte Quanteon and analyzed using FlowJo v10.2.

### Statistical analysis

Comparison of the four groups (naïve unvaccinated, infected unvaccinated, naïve vaccinated, infected vaccinated) based on previous infection and vaccination as well as different timepoints were investigated by one-way ANOVA and paired t tests using GraphPad Prism 9.3.1 (RRID: SCR_002798). In order to compare the slopes of the four groups, the antibody concentrations were log transformed, and the slopes after a linear regression analysis were compared by ANOVA. Statistical significance was denoted as *p<0.05, **p<0.01, ***p<0.001, and ****p<0.0001. p>0.05 was considered not significant (ns). Participants infected or vaccinated during the course of the study had their relevant timepoints included in multiple categories. For naïve unvaccinated participants, their changes in antibody titer were tracked starting with their earliest available timepoint. For infected unvaccinated participants, tracking started two weeks (0.5 months) after positive COVID nucleic acid amplification test (NAAT) or symptom onset; while for vaccinated participants (regardless of pre-immunity), tracking started two weeks (0.5 months) after receiving the complete primary series of vaccines to allow time for seroconversion. Completing the primary series is defined as having received two doses of the Pfizer-BioNTech or the Moderna vaccines or one dose of Johnson & Johnson’s Janssen vaccine.

## Results

In order to track and compare the changes in anti-RBD IgG antibody levels, the anti-RBD IgG antibody concentrations of 1,081 participants were tracked, based on a total of 3,970 individual timepoints (Figure 1, Table S1). Participants were broken up into four distinct categories: 1) The naïve, unvaccinated group (n=418) included participants who were never infected with the SARS-CoV-2 virus or vaccinated. 2) The infected, unvaccinated group (n=306) comprised of participants who had a confirmed SARS-CoV-2 infection either by NAAT, rapid antigen test, or a combination of COVID-19-specific symptoms followed by a corresponding significant rise in anti-RBD antibodies. 3) The naïve, vaccinated group (n=515) encompasses participants who were never infected with the SARS-CoV-2 virus but received their primary vaccine series. 4) The infected, vaccinated group (n=303) consists of participants who were pre-immune to the SARS-CoV-2 virus at the time of their vaccination and received their primary vaccine series.

**Figure 1:**
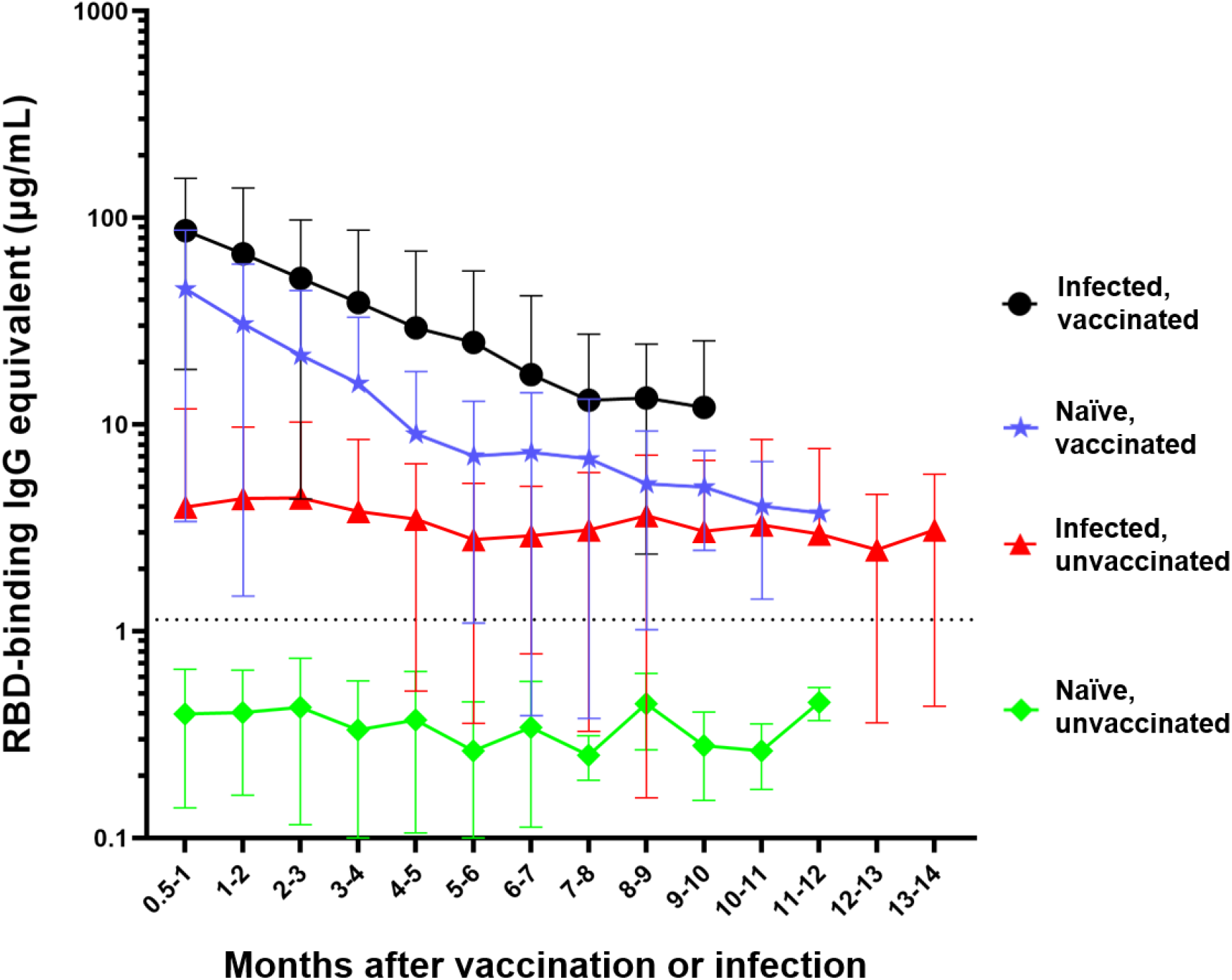
Differential waning of RBD-binding IgG antibody levels based on vaccination and infection status. Naïve unvaccinated (n=418) and infected unvaccinated (n=306) show no change in antibody levels over time (p>0.05); naïve, vaccinated participants (n=515) and infected, vaccinated participants (n=303) both show significant waning over the time (****p<0.0001). The antibody level of the naïve unvaccinated group was always lower than the other groups (****p<0.0001); the infected vaccinated group was always higher than any other group (*p<0.05); naïve, vaccinated group is higher than infected, unvaccinated group for the first 4 months after vaccination (**p<0.0014). The rate of decay was only significantly different between the vaccinated and infected groups (****p<0.0001) but not between the two vaccinated (p=0.7762) and the two unvaccinated groups (p=0.9476). Number of months start with the time of reception of the primary vaccines series for the vaccinated groups, time of infection for the infected unvaccinated group, and the first available timepoint for the naïve unvaccinated group.

The naïve, unvaccinated participants had antibody levels (mean = 0.4 µg/mL) that were below the experimentally determined concentration threshold and did not significantly change over time (p=0.19) (Figure 1)^10^. Participants who were infected but unvaccinated seroconverted with an average initial IgG antibody level of 4 µg/mL. There was no significant waning of antibody levels in these participants (p=0.3169) over the first 14 months following infection (mean concentration across all timepoints = 3.6 µg/mL). There was a much greater initial mean antibody level (44.8 µg/mL), in naïve vaccinated participants, however, this concentration significantly decreased (****p<0.0001) each month for the first 6 months. The infected vaccinated group initially seroconverted even higher (mean = 86 µg/mL), however, the antibody level experienced significant waning (****p<0.0001), each month for the first 5 months (Figure 1).

The anti-RBD IgG antibody concentrations of the naïve unvaccinated group were significantly lower than the antibody concentrations observed in any of the other three groups at all timepoints (****p<0.0001) (Figure 1). Participants who were infected and then vaccinated had significantly higher antibody levels than participants in the other three groups (***p<0.001 compared to infected unvaccinated and naïve unvaccinated groups, *p<0.05 compared to naïve vaccinated group). Naïve participants who were vaccinated had significantly higher anti-RBD antibody concentrations compared to unvaccinated participants who were previously infected for the first four months (****p<0.0001 for the first three months, **p=0.0014 during the fourth month) and no longer showed a significant difference beyond that timepoint (Figure 1). The slopes of each of the four groups were significantly different from every other group (****p<0.0001) with the exception of the two unvaccinated groups (p=0.9476) and the two vaccinated groups (p=0.7762).

Out of the 1,081 participants randomly selected for the longitudinal serum analysis, the antibody responses of participants who received two base mRNA vaccinations and an mRNA booster vaccine dose were assessed in 306 participants (Figure 2). Most of the participants received a homologous booster (227 Pfizer-BioNTech and 55 Moderna), but 24 received a heterologous booster vaccine dose (13 with Pfizer base + Moderna booster, 11 Moderna base + Pfizer booster). On average, participants experienced a 14-fold increase in their antibody levels compared to their last pre-boost timepoint (****p<0.0001), reaching a mean antibody concentration of 119.2 µg/mL (Figure 2). Moreover, their antibody levels were also on average 3 times higher than initially 2-4 weeks after receiving their second base vaccine dose (****p<0.0001), which was on average 39.8 µg/mL and remained significantly higher for the first 3 months (**p=0.0012). After the reception of the booster dose, significant waning was observed (****p<0.0001 between the first and second month, *p<0.05 between the second and third month), but the antibody levels were still significantly higher 5 months after the booster compared to the level immediately preceding the booster (**p=0.0067).

**Figure 2:**
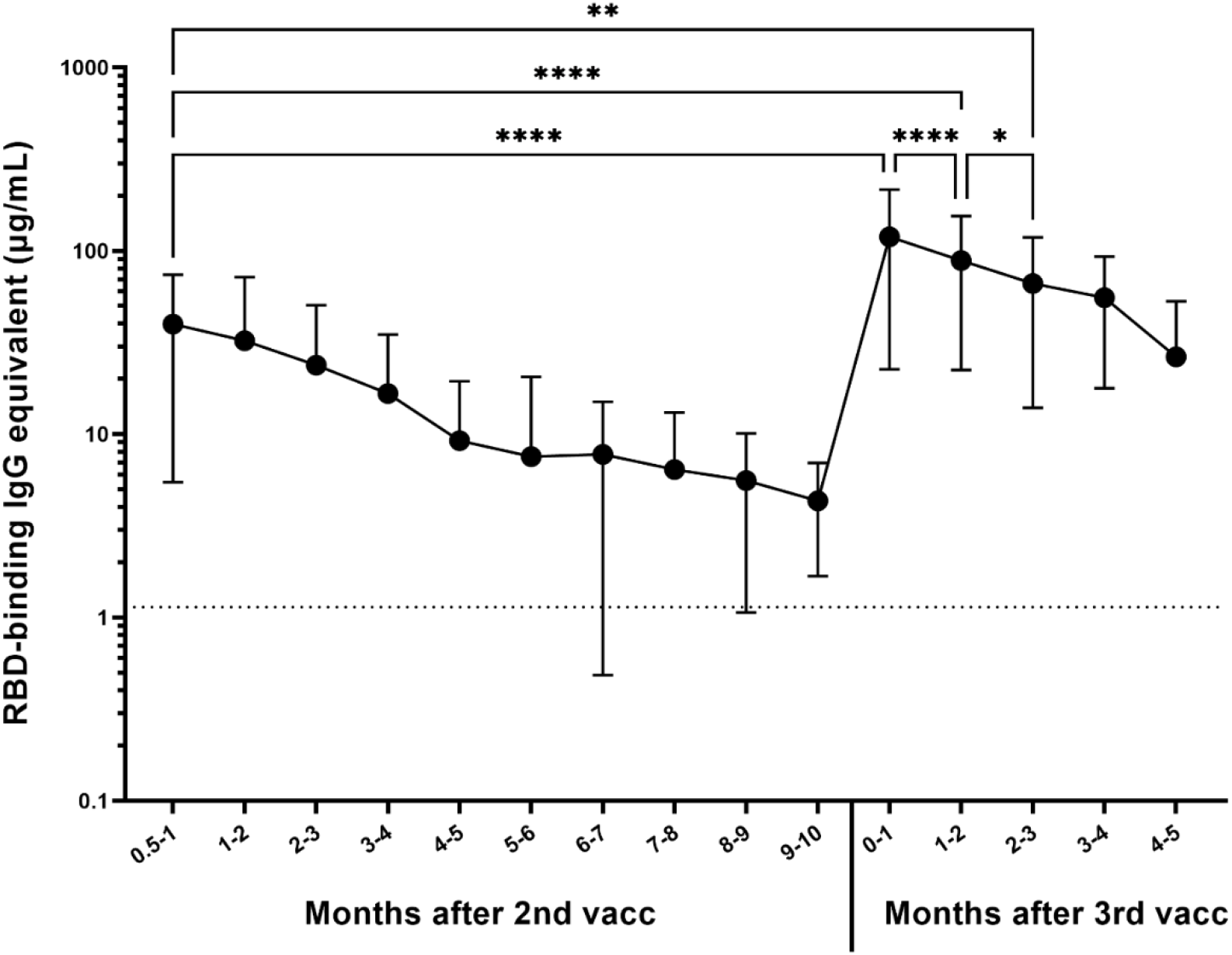
Response to 3^rd^ dose of mRNA vaccines (n=306). Significant increase in anti-RBD IgG antibody level was observed between the last available pre-booster timepoint and the timepoint within the first month after the booster (****p<0.0001). The difference between the timepoint 2-4 weeks after the second dose and the timepoint within the first month after the booster was also significant (****p<0.0001). Month to month antibody decay is significant for the first 3 months after the booster, at which point the antibody level was still significantly higher than 2-4 weeks after the second dose (**p=0.0012).

Out of the entire cohort, a total of 25 participants were identified who at some point during the study tested positive for RBD-binding IgG antibodies due to vaccination or infection, but have waned below the threshold into the negative range on a later timepoint (Figure 3A, Table S2). In order to confirm that these participant have in fact lost seroprotective status, viral neutralization tests using infectious SARS-CoV-2 USA-WA1/2020 strain were performed. Out of the 25 candidates, 12 either had non-detectable neutralization at the pre-waned timepoint (false positive by ELISA) or still had some level of neutralization potential on the post-waned timepoint (false negative by ELISA) (Figure 3B). The other 13 participants demonstrated some level of neutralization at their pre-waned timepoints but no neutralization at their post-waned timepoint, confirming the ELISA results that the participants have in fact lost their pre-existing seroprotection (Figure 3B). Serum collected from the post-waned timepoint from these 13 participants (10 post-vaccination and 3 post-infection) underwent *in vitro* differentiation, wherein the PBMCs were stimulated by recombinant IL-2 and R848 to induce a cellular memory recall response. In addition, two normal converters (CVI-004 and P-073) – participants who seroconverted after vaccination as expected – and four non-converters – participants who never experienced a significant increase in antibody levels after vaccination or infection – were also included as controls. The normal converters had a significant level of total IgG, Cal/09 IgG, RBD IgG, and spike IgG antibodies, while the non-converters only had a significant level of total IgG antibodies, signifying that the experiment was successful at eliciting a *de novo* antibody response (Table 1). Of the 10 participants participants who lost seroprotected status post-vaccination, 7 had a significant recall response against RBD and spike, while the other 3 did not (Table 1). Two of those three also showed no significant Cal/09 antibody levels. None of the 3 participants who lost seroprotected status post-infection demonstrated a significant recall response against RBD or spike, while 2/3 showed a significant Cal/09 response (Table 1).

**Table 1:**
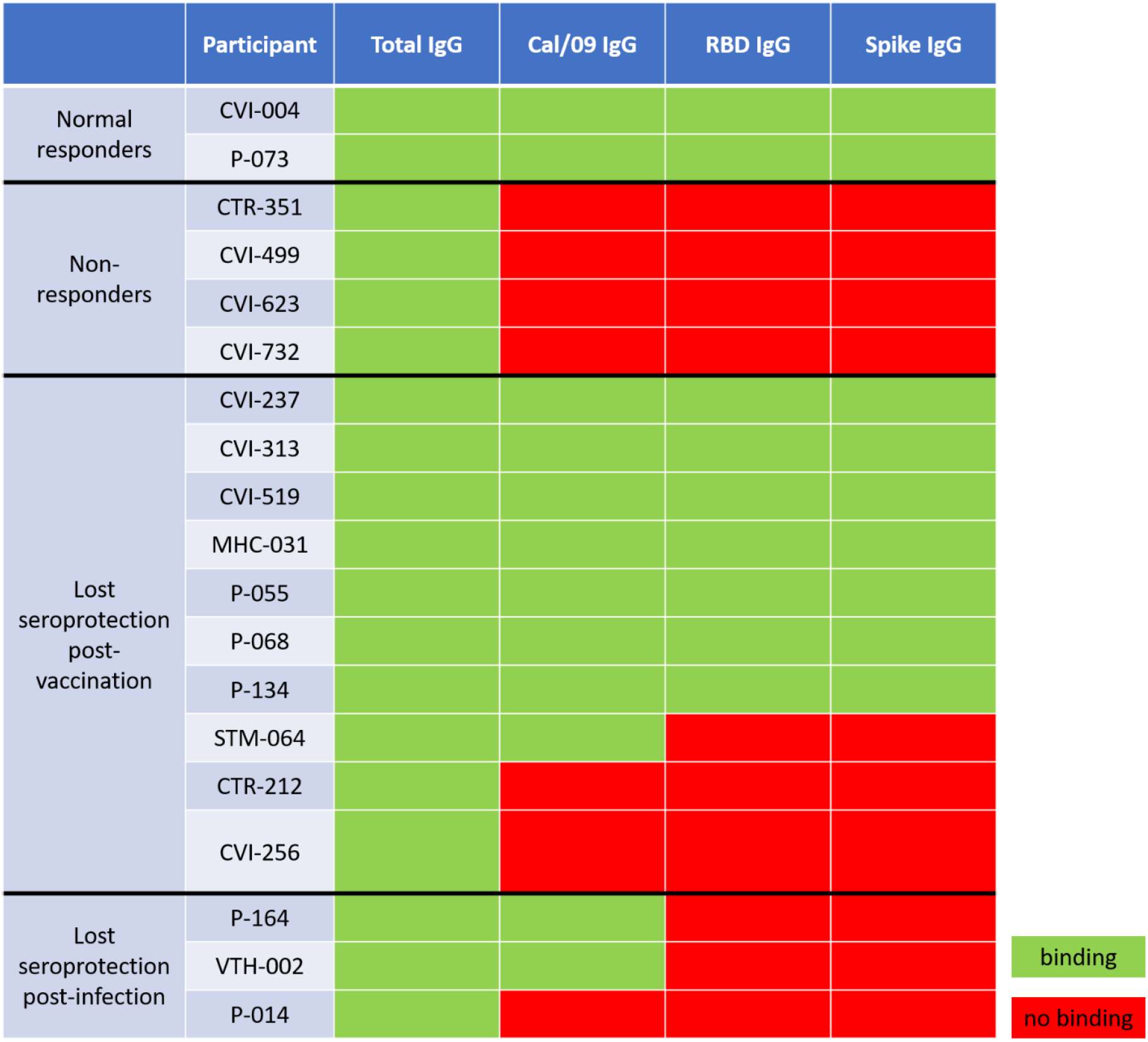
Memory B cell recall response in participants who lost their seropositive status after vaccination or infection. Differentiated PBMC supernatants from the post-waned timepoints were tested for total IgG antibody levels, as well as IgG antibodies specific to A/H1N1/California/2009 pandemic influenza strain. Normal responders seroconverted as expected, non-responders failed to seroconvert after vaccination, lost seroprotection groups initially seroconverted after vaccination or infection but later lost seroprotective status as shown by a lack of both RBD-binding and SARS-CoV-2 neutralizing antibodies.

**Figure 3:**
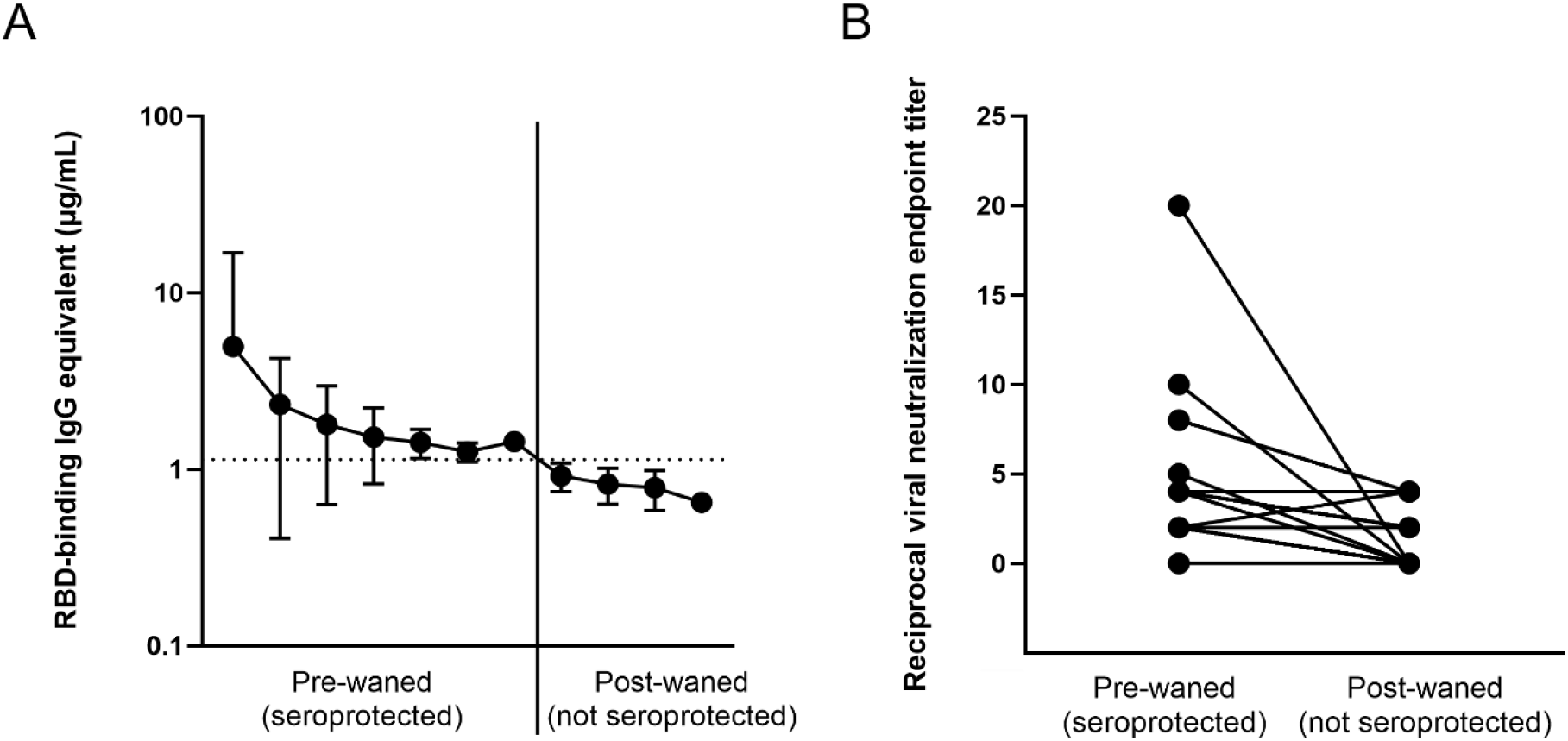
Loss of antibody protected status based on (A) antibody binding, and (B) viral neutralization in waned participants. All 25 participants demonstrated a loss of anti-RBD binding antibodies (A), but only 13 showed a loss of neutralization potential (B), signified by an initial non-0 neutralization endpoint titer and an endpoint titer of 0 at the post-waned timepoint.

Due to limited availability of PBMCs, B cell immunophenotyping could only be performed on 7 participants who waned post-vaccination (Table S3). 6 of the 7 were from participants who showed a significant SARS-CoV-2-specific memory recall, while the seventh (CVI-732) lacked all antigen-specific B cells after *in vitro* stimulation. While 20.4% of CD19^+^ CD20^+^ cells were memory B cells in the first 6 participants raging from 7% to 21% of IgG^+^ cells (median = 11.7%), CVI-732 demonstrated no detectable CD19^+^ B cells by flow cytometry (Table S3).

## Discussion

A few months after the World Health Organization declared the global COVID-19 pandemic in the spring of 2020^21^, participants were enrolled in the SPARTA program and tracked for immune responses elicited by SARS-CoV-2 infection. Longitudinal samples were collected from over 1,000 participants prior to and after infection and subsequently following vaccination. The data in this report analyzes the immune responses prior to the omicron variant wave that began in December 2021.

There was a stark difference in the waning of antibody levels in vaccinated versus infected participants. While infection-induced antibody levels were ∼10-fold lower than vaccinated participants, there was no significant waning over the next 14 months after infection. While vaccination-induced antibody levels measured 2-4 weeks post-vacination were significantly higher than infection-induced antibodies, they waned quickly, eventually asymptoting to levels similar to participants who were infected, but not vaccinated. Thus, the difference between the antibody levels in infected participants regardless of vaccination status is only apparent in the primary response and diminishes in the following months. Participants who were both infected and vaccinated had significantly higher antibody levels than infected unvaccinated participants for over 10 months. In contrast, participants who were never infected with SARS-CoV-2 before vaccination had significantly higher antibody levels compared to infected but unvaccinated participants for only the first 4 months following vaccination. The most robust antibody levels were reached and maintained by participants who were infected prior to vaccination. However, there was no significant difference between the rate of waning between the naïve vaccinated and the infected vaccinated groups, suggesting that pre-immunity does not significantly affect the rate of the waning of antibodies after vaccination. Thus, the differences between the two vaccinated groups were entirely due to the higher level of initial seroconversion amongst pre-immune participants. As the half-life of human IgG antibodies is 2-4 weeks, it is evident that the level of waning is proportionally related to the antibody level, and as their antibody levels dwindle, they asymptote instead of continuing to decline in a linear fashion^22,23^.

Antibody waning following mRNA vaccination has been previously reported^24–27^, and booster vaccine doses were approved in order to help to keep circulating antibody levels high. However, antibody waning may not be relevant, since a high level of circulating antibodies is not necessary for an effective B memory recall response^13,28,29^. In order to explore this dichotomy, this study assessed the effect of administering a 3^rd^ mRNA vaccine dose on antibody levels, and well as the B cell memory recall response in participants with undetectable levels of anti-SARS-CoV-2 antibody levels following initial presence of antibodies after vaccination.

Participants who received a booster vaccination had a 14-fold increase in anti-RBD antibody levels, and their post-booster antibody levels were 3 times higher than it was after receiving their second dose. This suggests that not only do boosters effectively increase the quantity of circulating serum antibodies, but they also generally provide more antibodies capable of binding to the RBD region of the spike protein. While there is still significant waning following the booster dose, antibody levels stay higher for a longer period of time.

Memory B cell recall occurs in the vast majority of SARS-CoV-2 vaccinated and convalescent individuals^30–33^. While the waning of antibody levels is natural due to the short half-life of antibodies^22,23^, concerns were raised about what happens to a very specific subset of participants who had a blunted serological response demonstrated by significantly lower initial antibody concentrations after vaccination (*p=0.026; mean for cohort = 44.8 µg/mL, mean for subset = 11.77 µg/mL). This subset of participants subsequently wane below the level of seroprotection^13^. In order to explore the effectiveness of B cell memory recall in the absence of detectable circulating antibodies, the SARS-CoV-2 spike-specific recall response in differentiated PBMCs from participants who lost their seroprotected antibody levels after vaccination or infection was assessed. Though the anti-RBD ELISA had a sensitivity and specificity of 96%^10^, most of these participants were close to the threshold and had a higher likelihood of being categorized as false positive or false negative. In order to verify that they did indeed possess protective antibodies, which later declined to undetectable levels, VN assays were performed in order to confirm seroprotected status. While analysis of binding antibody levels is useful and highly correlated with neutralization titers^10,34–36^, VN assays using naturally occurring infectious SARS-CoV-2 is useful to assess not only the ability of antibodies to bind a small portion of the virus but also to quantify all antibodies capable of preventing cytopathic effects elicited by the entire live virus, creating a more realistic *in vitro* model.

The 13 participants whose anti-RBD antibodies declined below the threshold of protection based on ELISA also lost their previous ability to neutralize virus (Table S2). In addition, two normal vaccine responders and four who never seroconverted after vaccination were included as controls. After *in vitro* PBMC stimulation with IL-2 and R848, all normal responders were able to recall SARS-CoV-2-specific antibodies, while the non-responders did not have a significant SARS-CoV-2 antigen-specific recall, but still produced non-specific IgG antibodies *de novo* (Table 1). Three of the four non-responders had autoimmune conditions or were taking immunosuppressant medication (Table S2). Of the 13 participants who lost previously existing seroprotected status, 10 initially seroconverted due to vaccination, while the other three initially seroconverted due to a SARS-CoV-2 infection. All three had asymptomatic infections. None of the 3 previously infected and only 7 out of 10 vaccinated participants showed a significant SARS-CoV-2-specific recall response. B cell immunophenotyping confirmed that 6 of the participants who had a robust memory recall had a significant number of IgG^+^ memory B cells (Table S3). On the other hand CVI-732, a non-responder taking the corticosteroid medication Prednisone^37^, as well as the anti-CD20 monoclonal therapy rituximab^38,39^ for rheumatoid arthritis, had no antigen-specific antibody recall post-stimulation and no detectable B cells by flow cytometry (Table 1, Table S2, Table S3). Future studies will focus on the effect immunosuppressants and autoimmune conditions have on serological and cellular responses.

There have been reports of blunted recall response after infection with coronaviruses in previous studies. One such study from 2011 detected no memory B cells in any of the 23 people who recovered from SARS-CoV infection, while a memory T cells response was present in more than half^40^. Another study found that SARS-CoV-2 may blunt the germinal center response due to a block in Bcl-6^+^ T_FH_ cell differentiation, an increase in T-bet^+^ T_H1_ cells, and aberrantly high TNF-α levels, causing a diminished recall response^41^. Our finding that antibody recall to Cal/09 was also absent in all non-responders and in some who lost seroprotection to SARS-CoV-2 suggests that while the overall IgG recall is substantial, the antigen-specific recall response in these participants is blunted not only against SARS-CoV-2 but also the Cal/09 pandemic influenza strain, perhaps due to overall lower level of total IgG antibodies (Table 1).

One of the limitations of this study is that we relied on accurate reporting of demographic information, comorbidities, previous SARS-CoV-2 testing and symptoms, and vaccine information by the participants. While the sample size for each of the four infected/vaccinated groups compared is robust, not all timepoints were available for all participants. Only participants who were convalescent prior to vaccination were reported in the infected vaccinated group, participants who had breakthrough infections after vaccination will be included in a future study. The sample size for the *in vitro* differentiation experiment was low due to the uncommon nature of losing seroprotective status amongst our participants. All participants who lost their confirmed seroprotected status were included in the analysis; future studies are necessary to strengthen our findings with additional participants.

In conclusion, vaccinated groups seroconverted to higher antibody levels and experienced significant antibody waning regardless of pre-immunity. In contrast, infected unvaccinated participants had no significant waning during the first 14 months after infection. The rate of antibody waning was not significantly different between pre-immune and immunologically naïve participants. Antibody levels are greatly increased by the administration of a booster dose, to a level 3-fold higher than their previous peak antibody level 2-4 weeks after the second dose. Memory B cell recall responses were absent in all infected and 30% of vaccinated participants who lost seroprotected status. Thus, the relationship between circulating anti-SARS-CoV-2 antibody levels and B cell memory recall may be more tightly linked than expected. Therefore, maintaining high circulating antibody levels by administering booster doses could be a highly effective way to counteract antibody waning and to avoid excessive waning of antibody levels.

## Acknowledgements

The authors would like to thank all participants enrolled in the SPARTA study. The authors would also like to thank the SPARTA collection and processing teams in Athens, Augusta, and Los Angeles, especially Brittany Baker, Charlotte Bolle, Debbie Bratt, Courtney Briggs, Jasmine Burris, Jordan Byrne, Patrick Fagan, Naveen Gokanapudi, Omar Hamwy, Tejal Hill, Lauren Howland, Hana Ji, Michael Kulik, Katie Mailloux, Patty Ray, Hua Shi, Cleopatria Smith, Donna Phan Tran, Terrie Waits, and Emma Whitesell,. We would like to thank Deborah Keys and Giuseppe A. Sautto for the immunological and statistical consulting. We acknowledge the staff at the Animal Health Research Center at the University of Georgia for the upkeep and maintenance of the BSL-3 facilities. The recombinant proteins were produced by Spencer Pierce, Ethan Cooper, and Jeffrey Ecker, and the team in the Center for Vaccines and Immunology protein production core. We also thank the entire staff at the University of Georgia Clinical and Translational Research Unit (CTRU) for assistance in collecting samples for the SPARTA program. The CTRU was supported by the National Center for Advancing Translational Sciences of the National Institutes of Health under Award Number UL1TR002378. The content is solely the responsibility of the authors and does not necessarily represent the official views of the NIH. Funding for this study was provided by by the National Institute of Allergy and Infectious Diseases (NIAID), a component of the U.S. National Institutes of Health (NIH), Department of Health and Human Services, under contract 75N93019C00052, and the University of Georgia (US) (UGA-001). In addition, TR is supported by the Georgia Research Alliance as an eminent scholar (GRA-001).

## Supplementary Information

**Table S1: List of participants whose antibody levels were longitudinally tracked**.

**Table S2: List of participants whose memory recall response was measured**.

**Table S3: Frequency and number of B cells indicated by direct *ex vivo* B cell immune cell phenotyping**.

**Figure S1: Memory B cell gating strategy**.

