## Supplementary figures and images for "The effect of waning on antibody levels and memory B cell recall following SARS-CoV-2 infection or vaccination"

### Figure S1

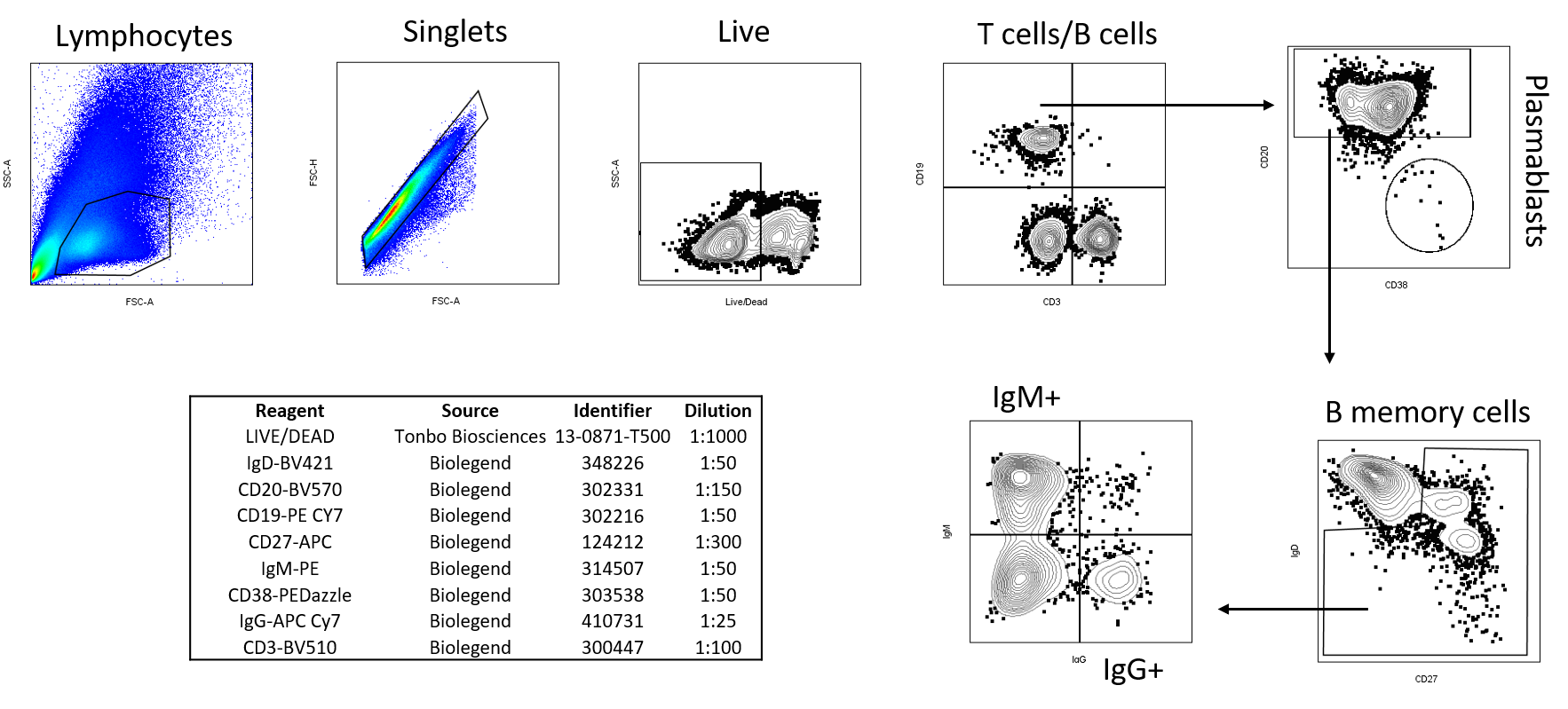
